# ADAR1 A-to-I RNA editing alters codon usage

**DOI:** 10.1101/268813

**Authors:** Pavla Brachova, Nehemiah S. Alvarez, Xiaoman Hong, Kailey A. Vincent, Keith E. Latham, Lane K. Christenson

## Abstract

**Background:** Fully grown mammalian oocytes and eggs are transcriptionally quiescent, and therefore have a unique RNA environment in which cellular processes depend on post-transcriptional regulation. RNA editing of adenosines into inosines (A-to-I) by adenosine deaminases acting on RNA (ADARs) is a common post-transcriptional gene regulatory mechanism, yet it has not been systematically studied in oocytes.

**Results:** A genome-wide RNA editing analysis of transcriptionally active growing oocytes from postnatal day 12 (PND12) mice, fully grown germinal vesicle (GV) oocytes, and transcriptionally quiescent metaphase II (MII) eggs indicates an abundant amount of A-to-I editing of mRNA transcripts. Editing of mRNA was greatest in GV oocyte and MII eggs compared to the PND12 immature oocytes, this was consistent with ADAR1 levels within these cells. Compared to somatic tissues, oocytes exhibited a different pattern of RNA editing, with a high proportion of RNA edits occurring in the coding regions. These edits resulted in nucleotide substitutions that were enriched at the third nucleotide of the codon (wobble position). Codon usage can affect mRNA stability and translation efficiency.

**Conclusions:** RNA editing in mouse oocytes is distinct from RNA editing in somatic cells due to increased frequencies of coding sequence RNA edits. We provide evidence in support of a previously unreported phenomenon of selective ADAR1 editing of the codon wobble position. Editing of the wobble position has the potential to fine tune post-transcriptional gene regulation through altering codon usage. This important observation advances our current understanding of RNA editing in mammalian cells.

## Background

The mammalian oocyte is a unique cell because it exhibits high transcriptional activity during growth, followed by transcriptional quiescence when it is stimulated to resume meiosis [1–3]. The oocyte then relies on a pool of previously transcribed and stored RNA to maintain cellular processes through fertilization and activation of the embryonic genome. Post-transcriptional gene regulation and translational control have increased importance in oocytes and are essential to generate high quality female gametes [4,5].

Comparison of RNA-seq datasets to genomic sequences has led to the realization that RNA editing is a widespread mechanism of post-transcriptional gene regulation [6]. RNA editing can affect both non-coding and coding regions of transcripts by altering splice acceptor sites, RNA secondary structure, recognition motifs in 5’ and 3’UTR, as well as codon sequences [7]. The result of RNA editing can impact transcript abundance, increase proteome diversity, and prevent detection of cellular RNA as foreign [8–16]. A common form of RNA editing is deamination of adenosines into inosines (A-to-I). This is carried out by adenosine deaminases acting on RNA (ADARs), a family of dsRNA binding proteins [17,18]. ADARs deaminate both inter-and intramolecular dsRNA of more than 20 base pairs in length [19], and are conserved in metazoans [20].

Mammals have three ADAR genes: *Adar*, *Adarb1*, and *Adarb2*, encoding the proteins ADAR1, ADAR2, and ADAR3, respectively [8]. ADAR1 and ADAR2 are enzymatically active and are known to edit dsRNA, whereas ADAR3 has no reported enzymatic activity. ADAR1 is ubiquitously expressed in tissues, while ADAR2 is expressed predominantly in the brain [21,22].

In addition to the three ADARs, another family of adenosine deaminases (testis nuclear RNA-binding protein TENR, also known as ADAD1), and TENR-like (TENRL, also known as ADAD2) have a similar structure to ADAR, but no deaminase activity due to the lack of a portion of the deaminase domain [23,24]. Deletion of ADAR1 in mice results in embryonically lethality due to defective organ development coupled with a systemic interferon response [12,25–28]. ADAR2 knockout mice die before postnatal day 20 due to unedited *Gria2* in neurons [29]. While ADAR2 has a defined molecular mechanism of action primarily in neurons, ADAR1 likely has a larger pool of edited substrates and a prominent role in post-transcriptional gene regulation in a variety of cell types.

Post-transcriptional events such as RNA editing are transient due to the rapid turnover of mRNA messages. Typically, in order to study post-transcriptional events in the absence of transcription, extra-physiological methodologies, such as chemical blockers or genetic manipulation, are necessary to dissect the molecular mechanisms [30–32]. To avoid this limitation, we studied RNA editing under a normal physiological condition in which transcription is naturally arrested. We utilized mouse oogenesis as model to study RNA editing due to the natural transcriptional quiescence that occurs during the later stages of oogenesis and maturation. In the current studies, we identified a previously unreported RNA editing phenomenon that has the potential to alter codon usage and affect mRNA stability and translation.

## Results

### ADAR expression and localization during mouse oogenesis

To dissect the role of RNA editing in mouse oocytes, we assessed transcript abundance of adenosine deaminases in growing PND12 oocytes, fully grown GV oocytes, and MII eggs. Globally, we observed that the three stages of oogenesis are transcriptionally distinct (Supplemental Fig 1a). We identified *Adar* (ADAR1) as the only adenosine deaminase with significant differential transcript abundance (Fig 1a, Supplemental Fig 1b-f). In mice, *Adar* generates three mRNA isoforms (Supplemental Fig 2). Transcript variant 3 (NM_001146296) encodes the longest ADAR1 protein, ADAR1 p150 (Supplemental Fig 2c). Transcript variant 1 (NM_019655) is similar to Variant 3, except for an alternative splicing event that shortens the 3’ end of exon 7 by 26 amino acids (Supplemental Fig 2b-c). ADAR1 p110 is encoded by transcript variant 2 (NM_001038587), generated from an alternative promoter within intron 1 (Supplemental Fig 2a). The remaining adenosine deaminases, *Adarb1* (ADAR2), *Adarb2* (ADAR3), and *adenosine deaminase domain containing 2 (Adad2)* exhibited low expression, while *Adad1* was undetectable (Supplementary Fig 1b-f).

**Figure 1.**
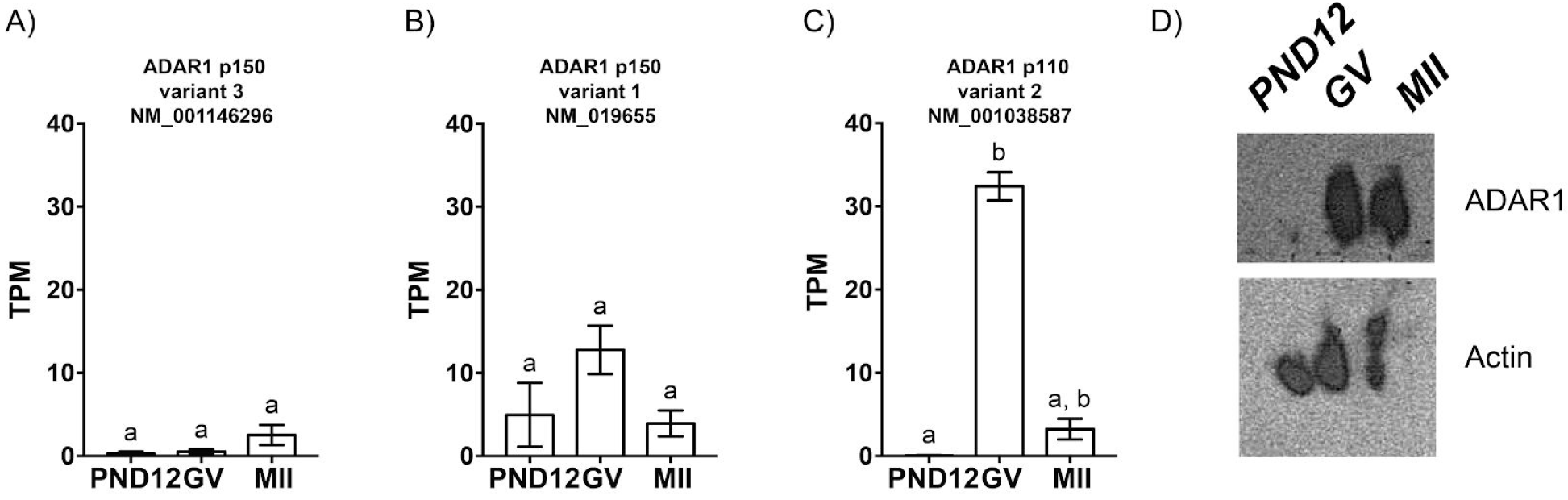
ADAR1 is predominantly expressed in mouse GV oocytes and MII eggs. Transcript abundance of *Adar* A) variant 3, B) variant 1, and C) variant2 in PND12 oocytes, GV oocytes, and MII eggs; TPM: transcripts per million. D) Western blot of ADAR1 and actin in PND12 oocytes, GV oocytes, and MII eggs; n=30 cells per lane. ^a,b^ Means ± SEM within a panel that have different superscripts were different (p < 0.05); Kruskal-Wallis test followed by Dunn’s multiple comparison tests.

Growing oocytes from PND12 mice expressed very low levels of transcripts encoding the ADAR1 p110 isoform (Fig 1a-c). Conversely, fully grown GV oocytes contained high levels of variant 2, which then decreased by the MII stage (Fig 1c). Neither of the transcripts encoding the larger ADAR1 p150 proteins showed statistical differences across the three stages of development (Fig 1a-b). Transcript variant 1 lacking a portion of exon 7 was present at higher levels in oocytes than transcript variant 3 (Fig 1b). Functional differences between transcript variant 1 and 3 are unknown, but the dsRNA binding domains and deaminase domain are identical (Supplemental Fig 2c). Abundant ADAR1 p110 protein levels were observed in GV oocytes and MII eggs (Fig 1d), with none detected in the growing PND12 oocytes. Additionally, we were unable to detect the p150 isoform of the protein (Fig 1d). These results indicate that *Adar*/ADAR1 is most abundant in transcriptionally inactive fully grown GV oocytes and MII eggs.

### Identification and validation of RNA edits in mouse oocytes

The differential expression of *Adar* in mouse oocytes and eggs led us to analyze global A-to-I editing during mouse oogenesis. We utilized the Genome Analysis Toolkit (GATK) pipeline for calling variants in RNA-seq data, with modifications [33]. We identified putative A-to-I RNA edits in pools of PND12 oocytes (n=3), GV oocytes (n=3), and MII eggs (n=7). Because growing PND12 oocytes had minimal levels of ADAR1, we expected minimal A-to-I editing to occur in these samples. Indeed, significantly fewer A-to-I RNA edited transcripts were observed in PND12 oocytes (733 ± 49 edited transcripts/sample; Mean ± SEM) than in GV oocytes (3,207 ± 117) and MII eggs (3,003 ± 290; p<0.05, one-way ANOVA; Fig 2a). RNA editing was validated in three identified genes (RPA1, MDC1, and WDR37) using Sanger sequencing of cDNA and genomic DNA derived from GV oocytes of wild-type C57BL/6J female mice (Fig 2b). We were able to successfully verify RNA editing in all three genes, as indicated by the appearance of A-to-G transitions on the forward cDNA strand, or C-to-T transitions on the reverse cDNA strand (Fig 2b).

**Figure 2.**
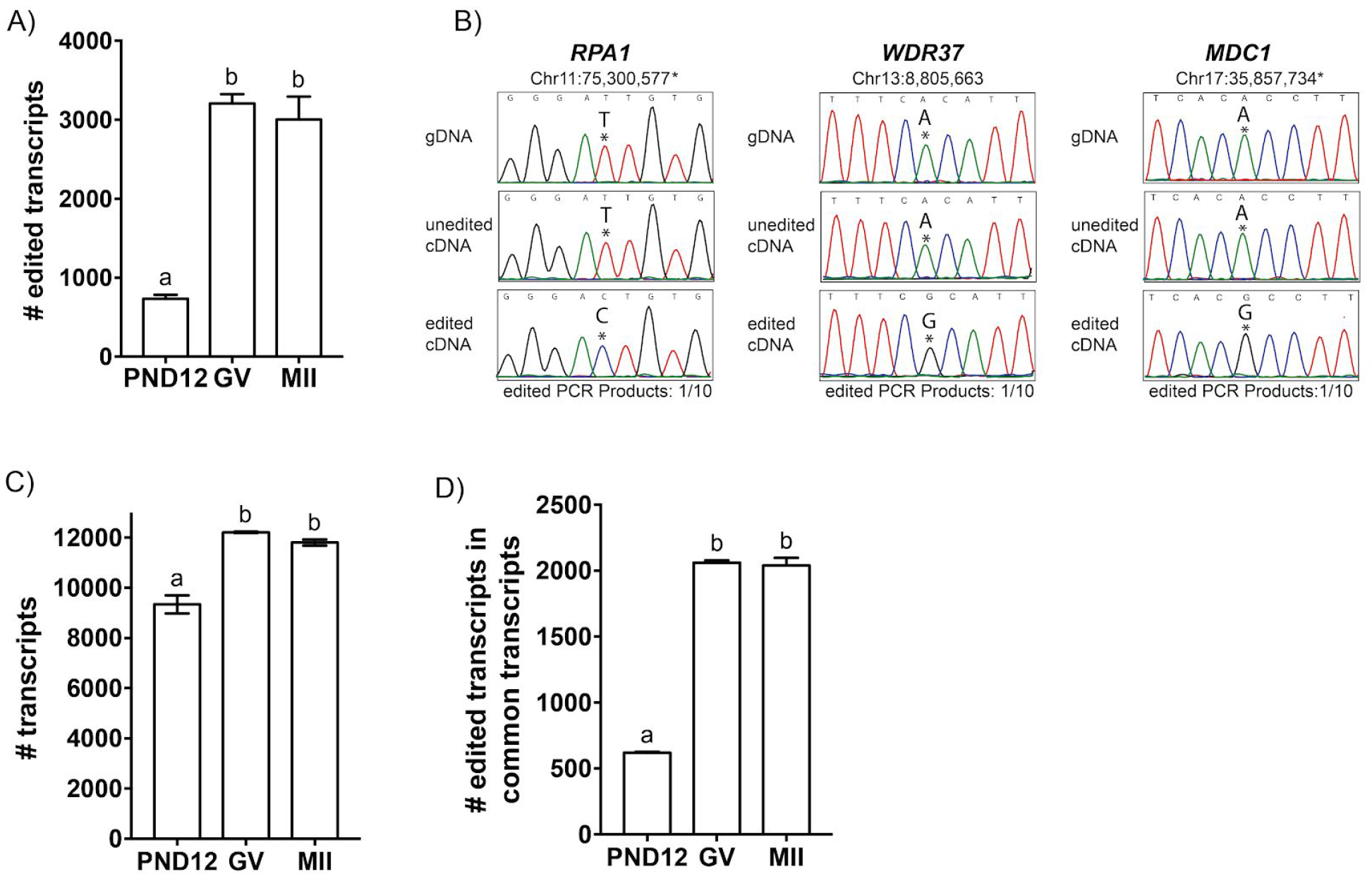
A-to-I RNA edits predominate in fully grown GV oocytes and MII eggs. A) The number of unique edited transcripts identified in PND12 oocytes, GV oocytes, and MII eggs. B) RNA edits were validated using Sanger sequencing of three genes. Chromosome location is indicated, and the * denotes minus strand of the DNA. C) The total number of mRNA transcripts per sample. D) The number of edited transcripts among commonly detected transcripts. ^a,b^ Means ± SEM within a panel that have different superscripts were different (p < 0.05); significance was determined using one-way ANOVA, followed by Tukey’s multiple comparison tests. Only transcripts with TPM ≥ 1 were analyzed.

It is possible that the ~4-fold increase in number of edited transcripts seen in GV oocytes and MII eggs represents a stabilized population of transcripts that are not present in PND12 oocytes, rather than due to ADAR1 activity. To test this, we measured transcript abundance between PND12 oocytes, GV oocytes, and MII eggs. We measured transcript abundance by aligning raw RNA-seq reads to the mouse RefSeq database which includes over 30,000 unique transcripts (See Methods). We observed that the total number of unique transcripts detected in GV oocytes and MII eggs was 30% greater than in PND12 oocytes (Fig 2c). An assessment of RNA editing in transcripts that were commonly present at all three stages of oogenesis (Fig 2d) was completed to account for increased transcript abundance observed in the later stage oocytes. Within the pool of common transcripts, we observed a ~4-fold increase in the number of edited transcripts in GV oocytes and MII eggs (Fig 2d). These results indicate that the increase in RNA editing observed in GV oocytes and MII eggs is not due to the overall amount of transcripts present, but is strongly correlated with oocyte ADAR1 abundance (Fig 1c-d).

### Oocyte-specific pattern of RNA editing

In addition to having the fewest edited transcripts, PND12 oocytes had the lowest proportion of the transcriptome that was edited (7.8 ± 0.2%; Mean ± SEM), compared to GV oocytes (26.3 ± 0.9%) and MII eggs (25.6 ± 2.6%; Fig 3a). The proportion of edited transcripts in oocytes was similar to mouse somatic tissues (colon, heart, large intestine, and stomach) from wild-type mice, except for the brain, which has a higher percentage of A-to-I edits (Supplemental Fig 3a). Consistent with higher levels of ADAR1 protein, GV oocytes and MII eggs have an approximately 2-fold increase in the number of RNA edits per transcript compared to PND12 oocytes (Fig 3b). We did not observe a difference in the number of edits per transcript among somatic tissues that express other ADAR genes (Supplemental Fig 3b and Supplemental Fig 4).

**Figure 3.**
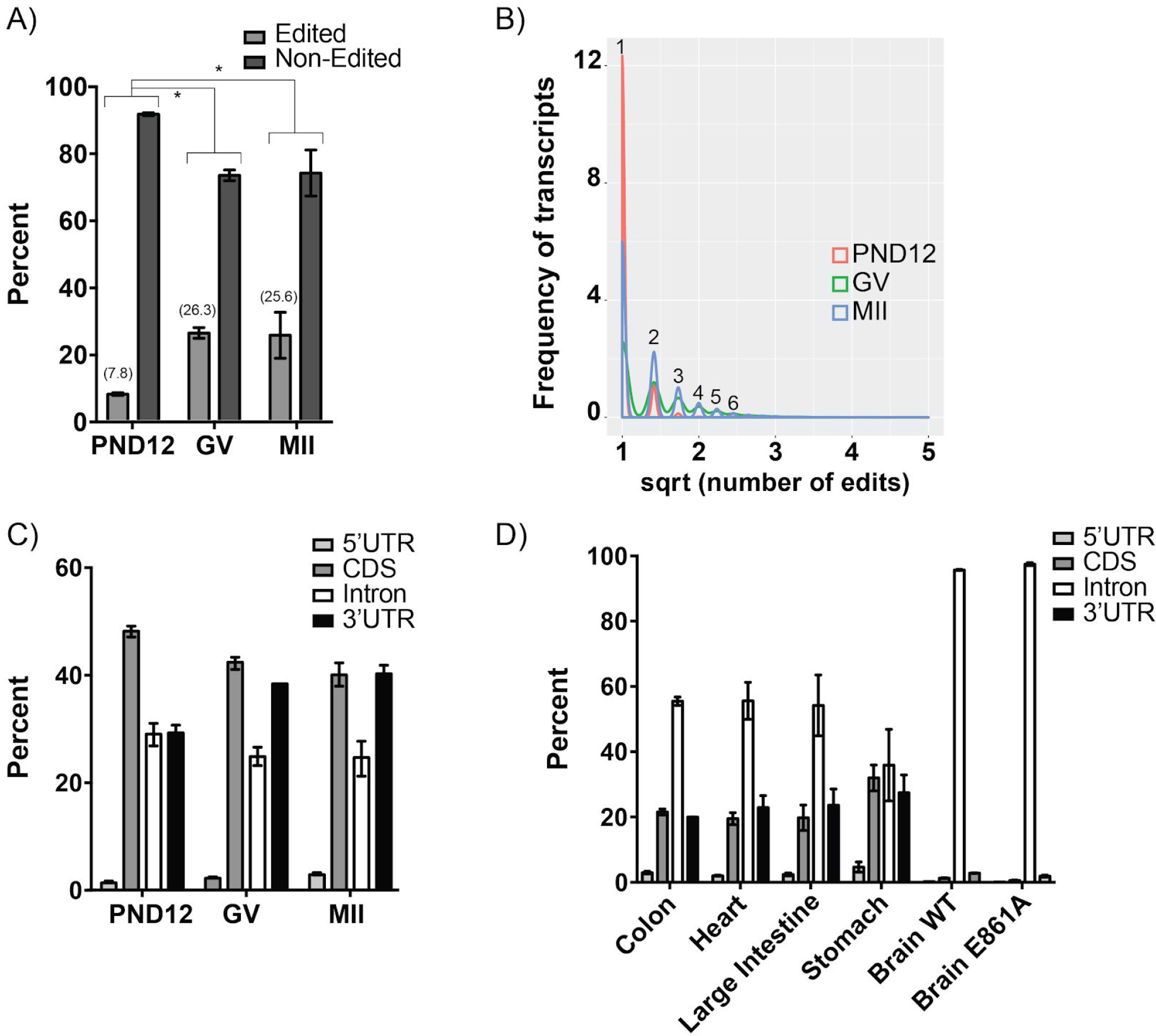
Distinct patterns of RNA editing in oocytes compared to somatic cells. A) Proportion of the transcriptome (percentage shown) that contains RNA edits within PND12 oocytes, GV oocytes, and MII eggs. B) The frequency of edited transcripts (y-axis) exhibiting one or multiple edits per transcript in PND12 oocytes, GV oocytes, and MII eggs. The x-axis is sqrt (square root) normalized, while the numbers above peaks represent the actual number of edits per transcript. Distribution of RNA edits within specific regions (5’UTR, CDS, intron, and 3’UTR) of mRNA of oocytes (C) and somatic tissues (D). *Means ± SEM within panel A are different (p < 0.05); significance was determined using X^2^ tests. Only transcripts with TPM ≥ 1 were analyzed.

To further characterize the nature of RNA edits in oocytes, we determined the location of edits within protein-coding genes by annotating the location of each edit (5’UTR, coding, intron, and 3’UTR). Overall, the distribution of edits is similar between all stages of oogenesis, however, compared to somatic tissue, the oocyte has a unique pattern of RNA editing (Fig 3c and Fig 3d). Mouse oocytes display a higher proportion of coding region (CDS) edits compared to mouse somatic tissue (Fig 3c and Fig 3d), indicating that RNA editing in mouse oocytes may be fundamentally different.

### Consequences of coding sequence RNA edits in mouse oocytes

To understand the consequence of RNA editing, if any, on the protein coding capacity of edited GV and MII mRNA transcripts, we used Ensembl Variant Effect Predictor (VEP) [34] to identify the synonymous and nonsynonymous substitutions (Fig 4a). Among the nonsynonymous substitutions, altered stop codons (stop loss, stop gain, or stop retained) made up less than 0.3% of coding sequence edits. Therefore, stop codon substitutions were not included in further analyses. We found a significant increase in synonymous substitutions in GV oocytes and MII eggs, coinciding with an increase in ADAR1 protein (Fig 4a and Fig 1d). Somatic tissues displayed a similar pattern to GV oocytes and MII eggs, which also express ADAR1 (Fig 4b). To test if the increase in synonymous substitutions was a result of increased ADAR1 activity, we utilized RNA-seq data from a genetic mouse model expressing a catalytically inactive ADAR1, with a point mutation at E861A [35]. The prevalence of synonymous substitutions dramatically decreased in the E861A mutant, similar to PND12 oocytes, which also do not express ADAR1 (Fig 4b and Fig 1d).

**Figure 4.**
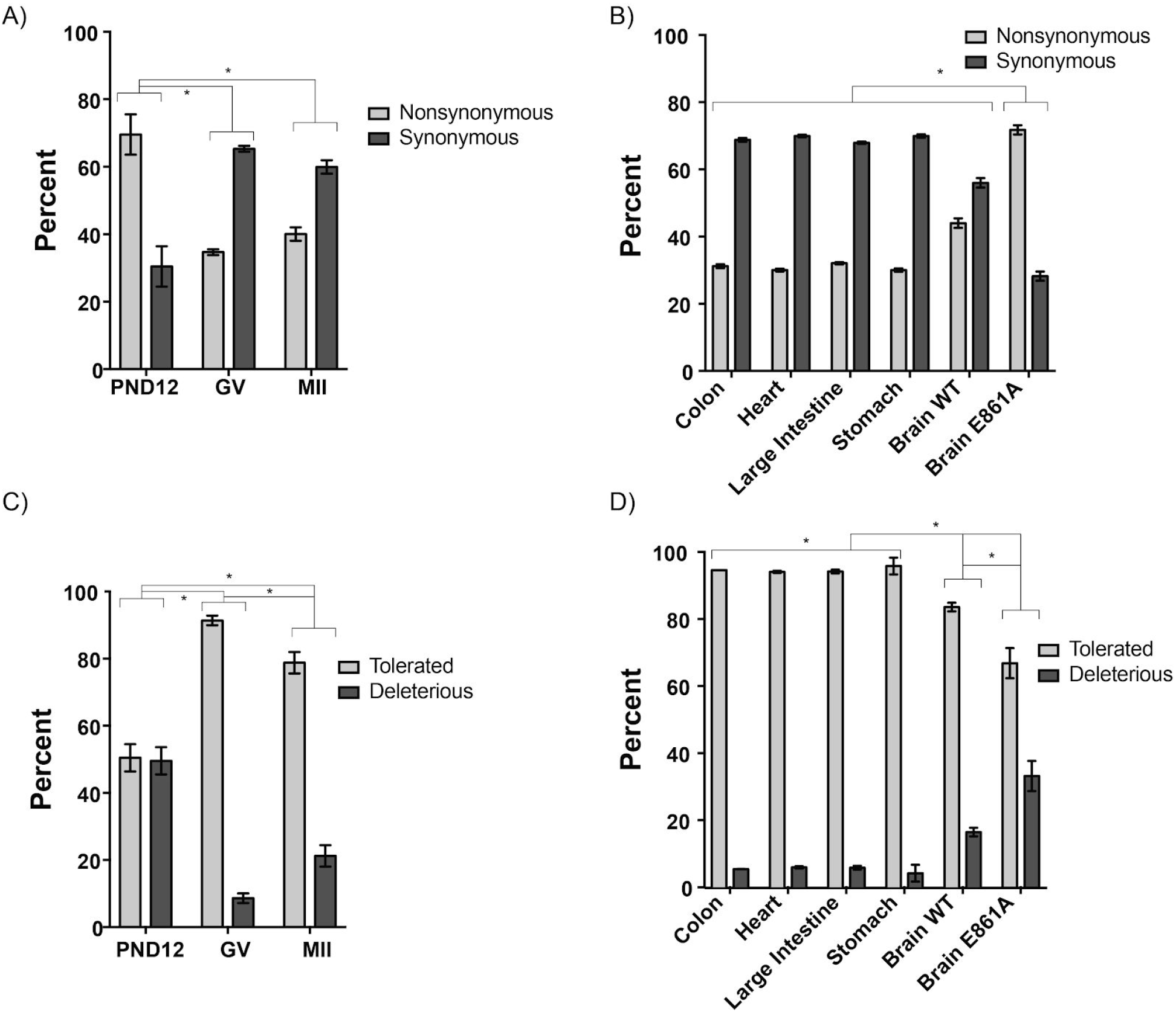
The consequence of RNA edits in CDS of mouse oocyte transcripts. A) The proportion of RNA edits that result in nonsynonymous or synonymous substitutions was determined for all edited mRNA transcripts in PND12 oocytes, GV oocytes, and MII eggs. B) The proportion of RNA edits in a variety of somatic tissues of wild type mice, as well as the ADAR1 E861A mutant brain tissue. The proportion of tolerated and deleterious transcripts following Sorting Intolerant From Tolerant (SIFT) analysis of the edited mRNA transcripts from oocytes (C) and somatic tissues (D). *Means ± SEM within a panel are different (p < 0.05); significance was determined using X^2^ tests. Only transcripts with TPM ≥ 1 were analyzed.

In order to predict the consequence of the amino acid substitutions, a computational tool, Sorting Intolerant From Tolerant (SIFT) that predicts the effects of amino acid substitution on protein function was used [36]. SIFT analysis showed that RNA edits observed in GV oocytes and MII eggs had increased levels of tolerated amino acid substitutions when compared to the PND12 oocytes (Fig 4c). Similarly, somatic tissues exhibited a much greater level of tolerated (>80%) versus deleterious substitutions (Fig 4d). In summary, ADAR1 editing is associated with increased tolerated, synonymous substitutions.

### Selective editing at the wobble codon position

The abundance of RNA edit synonymous substitutions led us to investigate the potential effects of ADAR1 editing on codon usage. We first determined the number of A-to-I edits that occur in the 34 different codons that contain an adenosine, excluding stop codons. We found that in GV oocytes and MII eggs, RNA edits occur more frequently in the following codons: AAA, ACA, CAA, CCA, GAA, and GCA (Fig 5a; p<0.05, two-way ANOVA). This effect was not due to a difference in the overall abundance of these codons (Supplemental Fig 5). Further analysis indicated that adenosine deamination appeared to be specifically enriched at the wobble position (Fig 5b). Even the AAA codon had a preference for RNA editing at the wobble position (Fig 5b). This occurred predominantly in GV oocytes and MII eggs, rather than in PND12 oocytes, consistent with the lack of detectable ADAR1 in PND12 oocytes (Fig 5b and Fig 1d). We also examined this phenomenon in all 34 codons containing adenosines, and observed a similar pattern of increased RNA editing occurring at the wobble position (Fig 5c) and in somatic tissues (Fig 5d). To test if RNA editing at the wobble position was due to ADAR1 catalytic activity, we compared brain tissues from wild-type mice with those of the ADAR1 E861A mutant. The ADAR1 E861A mutant appears to have a similar profile as PND12 oocytes, which also lack ADAR1 protein (Fig 5d and Fig 1d). Overall, ADAR1 activity was strongly associated with an RNA editing preference at the wobble position and this represents a novel ADAR1 signature.

**Figure 5.**
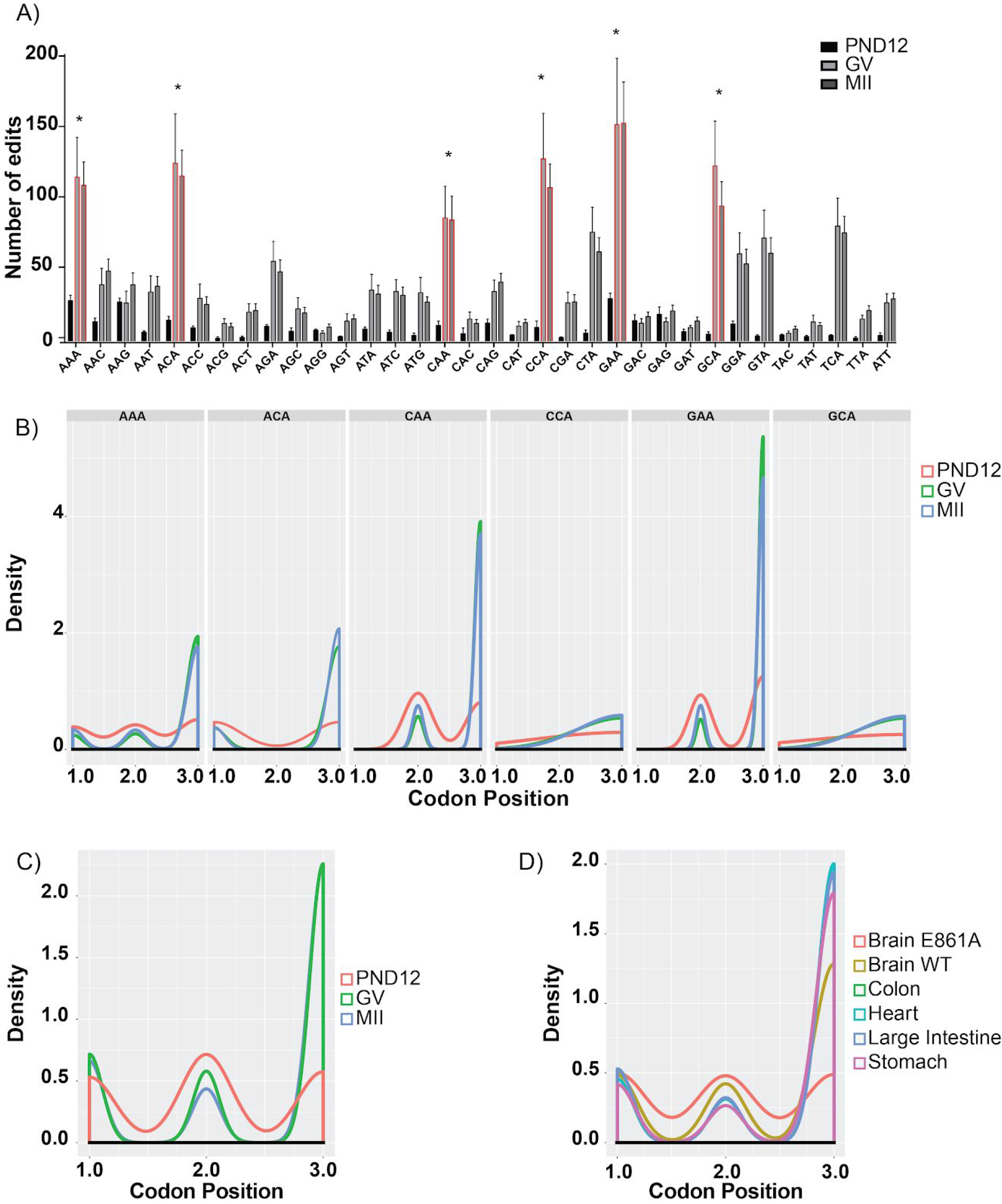
ADAR1 RNA editing activity is enriched at the wobble position of codons. A) Frequency of adenosine-containing codons in PND12, GV, and MII samples. B) Frequency of aRNA edit occurring at the first, second, or third codon position in PND12, GV, and MII samples. Global frequency of RNA edits occurring at the first, second, or third codon position in PND12, GV, and MII samples (C) and in somatic tissues (D). Means ± SEM within a panel that have superscripts ^*^were different (p < 0.05); significance was determined using two-way ANOVA tests. Only codons from transcripts with TPM ≥ 1 were analyzed.

### RNA editing of codons and transcript stability

It has been demonstrated that codon usage can affect mRNA stability [32]. We examined the potential consequences of ADAR1 codon editing on mRNA stability. In order to gain insight into how specific codons might affect mRNA stability, we relied on observations that demonstrate GV oocytes and MII eggs are transcriptionally inactive [3,37,38]. Therefore, changes in transcript levels during the transition from GV oocytes to MII eggs are due to RNA stability rather than *de novo* RNA synthesis. Differential RNA abundance calculations during the transition from GV to MII can be interpreted as RNA stability measurements.

Differential RNA abundance within GV oocytes and MII eggs was determined, and 477 transcripts were stabilized and 628 transcripts were destabilized (Fig 6a). From these transcripts, we determined the codon occurrence to mRNA stability correlation coefficient (CSC; [32] Fig 6b). We found that six codons frequently edited by ADAR1 were associated with stable mRNA (Fig 5a and Fig 6b). Moreover, editing within the codon would switch a stabilizing codon into a destabilizing one in three of these codons (Fig 6a). For example, the most frequently ADAR1 edited codon is GAA (Fig 6a), which is highly correlated with stable transcripts. If a GAA is hypothetically edited into GAG, it is then correlated with unstable transcripts (Fig 6b). ADAR1 RNA editing could affect overall mRNA stability through editing of codons.

**Figure 6.**
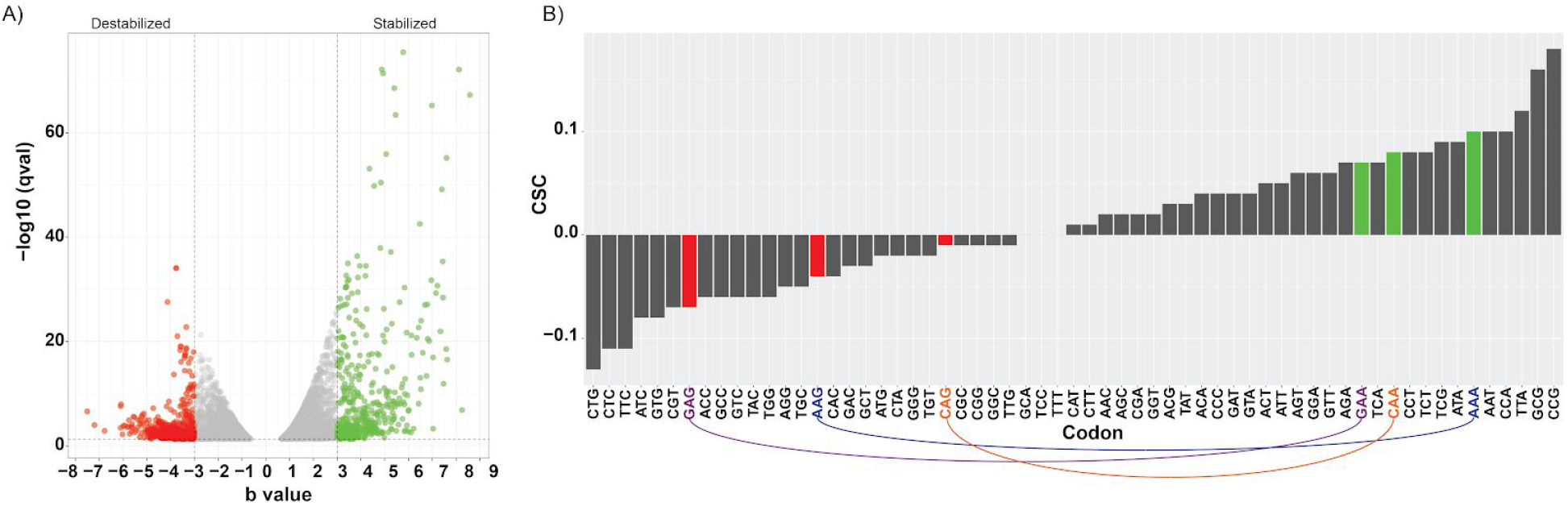
ADAR1 RNA editing destabilizes codons in mRNA. A) Volcano plot showing transcripts destabilized (red) or stabilized (green) when comparing mRNA from GV to MII. X-axis is denoted as a “b value”, an estimator of fold change from Sleuth. B) Codon stability coefficient (CSC) for each codon in the stabilized and destabilized fractions of transcripts. As a codon is hypothetically edited, lines link the stability of the unedited version to the edited version.

## Discussion

Our studies provide a genome-wide analysis of the post-transcriptional modification, A-to-I RNA editing, at different stages of mouse oogenesis and maturation. We determined that ADAR1 codon editing exhibited preferential modification of the wobble position. Codons with ADAR1 edits at the wobble position were correlated with unstable mRNA. Analysis of our data demonstrates the presence of a novel regulatory process involving RNA modifications during oogenesis that have the potential to regulate translation and mRNA degradation.

During the growth phase of oogenesis, mammalian oocytes accumulate and store maternal RNA that are required to support the subsequent transcriptionally silent stages of meiotic maturation, fertilization, and early embryonic growth prior to embryonic genome activation [1–3,38–40]. In fully grown GV oocytes and MII eggs, previously synthesized maternal mRNA are recruited into polysomes for translation [5,41]. Maternal transcripts are subjected to translationally coupled degradation, and in the absence of degradation, maternal transcripts accumulate, resulting in embryonic arrest [5]. Thus, the regulatory roles of post-transcriptional and translational control at this stage of oogenesis are essential to ensure successful embryonic development. We observed very little *Adar* expression and no detectable protein in transcriptionally active PND12 oocytes. This correlated with a reduced overall number of edited transcripts, decreased proportion of edited transcriptome, and a reduced number of edits per transcript. In contrast, fully grown GV oocytes and transcriptionally quiescent MII eggs expressed abundant ADAR1 protein and have increased levels of mRNA editing.

A distinct RNA editing pattern was observed in mouse oocytes, with a majority of A-to-I edits occurring in the CDS and 3’UTR regions of transcripts. This is unique from a variety of other mouse somatic cell types (Fig 3c-d; [44]), however, it is not unprecedented. Cephalopods have a high proportion of A-to-I RNA edits occurring in the CDS region of genes, and this has been hypothesized to increase proteome diversity and allow for increased adaptability [45]. A study of human oocytes identified a majority of A-to-I edits occurring in the 3’UTR regions (47.12 %), followed by intronic (33.77 %), non-coding RNA (17.12 %), and 5’-UTR regions (1.01 %), with only 0.98 % occurring in coding regions [46]. However, it is unclear if the increased coding region edits we have observed are due to species differences or due to the vitrification process and culture of human oocytes prior to sequencing, as described in the manuscript source of the oocyte sequencing data [47]. A recent cross-species analysis of RNA editing in somatic tissue revealed that species, rather than the type of tissue, was a greater source of RNA editing differences [6]. However, differences between oocytes and somatic post-transcriptional regulatory mechanisms are not unprecedented. For example, mouse oocytes express a unique isoform of Dicer, which favors endogenous siRNA biogenesis over miRNA biogenesis, that regulates the expression of retroelements [48]. It is possible that oocytes employ a unique A-to-I editing program necessary for steps in early embryonic development.

To understand the possible consequences of A-to-I editing within CDS regions of oocytes, we examined whether these edits cause nonsynonymous or synonymous substitutions within the mRNA. Our results indicate that GV oocytes and MII eggs, which have high levels of ADAR1 protein and A-to-I editing, have an increase in number of synonymous substitutions. Synonymous substitutions are known to affect RNA transcript stability as well as translation efficiency because of codon availability and codon bias [49]. Of the codons with adenosines that have the potential to be edited, we observed that GV oocytes and MII eggs contained six codons that were edited at a significantly higher frequency than PND12 oocytes (Fig 5a). These codons have a specific enrichment of RNA editing at the wobble position of the codon (Fig 5b). It should be noted that the increased appearance of certain edited codons was not a reflection of an overall increase in that codon within the mRNA transcripts present in oocytes (Supplemental Fig 5). To test if the preferential RNA editing of the wobble position was dependent on ADAR1 activity, we compared the editing profiles of brain tissue from wild-type mice to those of mice expressing an ADAR1 E861A mutation that eliminates enzymatic activity [50]. We observed that mice expressing the ADAR1 E861A mutation displayed significantly reduced editing at the wobble position. This indicates that the wobble position A-to-I substitution is regulated by ADAR1.

No current models of ADAR1 function explain the enrichment of A-to-I editing at the wobble position; however, it is not unprecedented that RNA editing can occur at specific codon positions. Mitochondrial genes of the slime mold *Physarum polycephalum* also have an increase in RNA edits occurring at the third codon position [51,52]. *P. polycephalum* mitochondrial RNA undergoes editing in the form of cytidine insertions at the third position of codons. In another example of RNA editing affecting codon usage, *Arabidopsis* and other plants exhibit deaminated cytidine bases primarily in the first two codon positions [53–55]. Although both of these examples involve a different type of RNA editing than ADAR A-to-I editing, they provide a antecedence for a link between RNA editing and codon position preference. *In vitro* models using purified ADAR1 and synthetic RNA have demonstrated that ADAR1 has a trinucleotide preference and specifically edits the second nucleotide of the triplet [56]. However, our analysis of codon editing using *in vivo* genetic models of catalytically inactive ADAR1 demonstrate that within the CDS, ADAR1 frequently edits the wobble position of codons. Most studies of ADAR editing do not focus on the CDS, and therefore this bias for editing the wobble position has gone undetected. Overall, our analysis produces RNA editing data that is consistent with previously reported editing profiles in somatic tissue. It is possible that mRNA from transcriptionally quiescent oocytes is edited at the first and second position of a codon, but degraded rapidly, resulting in an increase of mRNA with edits at the wobble position. However, even in transcriptional active somatic cells we observe that ADAR1 edits occur more frequently at the wobble position. Only in the catalytic ADAR1 E816A mutant and PND12 oocytes that lack ADAR1 editing, do we observe a decrease in wobble position editing. Further experiment are needed to understand the molecular mechanisms governing selective editing of codon positions.

## Conclusion

We identified extensive RNA editing within the coding region of genes and low intronic RNA editing in oocytes, contrasting with previously reported RNA editing profiles of mouse somatic tissues [44] and human oocytes [57]. Our study also demonstrates a previously unreported phenomenon of ADAR1 editing of the codon wobble position. The wobble position was edited at a higher frequency than any other position, and this was not an oocyte-specific phenomenon. We hypothesize that a consequence of ADAR1 editing is the fine-tuning of codon optimization in all tissue types. We therefore suggest that RNA editing by ADAR1 can contribute to mRNA stability through altered codon usage.

## Methods

### GV oocyte and MII egg RNA-seq datasets used in this study

Wild-type GV oocytes and MII egg RNA-seq data was identified by searching the Gene Expression Omnibus and downloaded from the Short Read Archive using SRAdb [58]. Wild-type GV oocyte data was used from data set SRP057558 [59]. The authors of this study demonstrated high quality of isolated GV oocytes (fully grown, meiotic competence, successful nuclear envelope breakdown and polar body extrusion). Contemporary GV oocytes were used for RNA-seq [59]. Wild-type MII oocyte RNA-seq data was used from data sets SRP034543 [60] and SRP065556 [61]. These samples were used because the MII eggs were isolated *in vivo*, from oviducts. Control somatic tissue RNA-Seq data was obtained from SRP098702 [35].

### Library preparation and RNA sequencing of postnatal day 12 oocytes

Three biological replicates from post-natal day 12 oocytes (6 oocytes per mouse, n=3) were isolated from the ovaries of wild-type C57BL/6J female animals. Ovaries were incubated with collagenase I (0.1% v/v, Worthington Biochemical Corporation, Lakewood, NJ) for 30 minutes to dissociate oocytes. During incubation with collagenase, the ovaries were pipetted up and down every 10 minutes to facilitate dissociation, and then washed through several droplets of culture medium without collagenase. RNA was extracted using the PicoPure RNA Isolation Kit (Thermo Fisher, KIT0204). RNA-seq libraries were prepared following the Nugen Ovation Ultralow Library Systems Protocol (Nugen®, Ovation® Ultralow Library Systems User Guide, M01219 v6). Briefly, first strand and second strand cDNA synthesis was conducted and purified using Agencourt RNAClean XP beads (Beckman Coulter, A63987). cDNA products were then amplified using SPIA amplification and purified using Qiagen MinElute Reaction Cleanup Kit (Cat# 28204). Amplified cDNA was then fragmented using a Covaris S2 Ultrasonicator, and digested using S1 Nuclease treatment. DNA was purified, end repaired, ligated to adaptors, and amplified. The resulting libraries were analyzed on a Bioanalyzer DNA Chip 1000. Fragment distribution was between 150–200 base pairs. RNA-seq was performed using single-end Illumina HiSeq 4000. This raw data has been deposited in SRA BioProject PRJNA434598, SRP133083.

### Transcript abundance analysis

Sequence alignment and transcript abundance calculations for PND12 oocytes, GV oocyte, MII eggs, and mouse somatic tissues were performed by aligning raw RNA-seq reads against *mus musculus* (mm10) transcripts curated from RefSeq using Kallisto, and differential transcript abundance was calculated using Sleuth [62–64]. Commonly edited and expressed transcripts were identified by performing multiple comparisons between all the samples (PND12 = 3, GV = 3, MII = 7).

### Identification of RNA edits and RNA edit consequence analysis

Raw RNA-seq reads were mapped to the *mus musculus* reference genome, mm10, using HISAT2 [65]. RNA/DNA differences were called following the Genome Analysis Toolkit (GATK) RNA-Seq variant pipeline with modifications [33]. The program elprep was used to sort, mark duplicates, and index RNA-seq reads [66]. Known SNPs were filtered out using a database from 17 mouse strains maintained by dbSNP [67]. The resulting VCF files were filtered for A/G and T/C variants. Variants calls were stranded with A/G occuring on the sense strand and T/C occuring on the antisense strand [68]. VCF files were used as input for Ensembl Variant Effect Predictor (VEP) [34]. VEP was used to identify edited transcripts, categorize the location within the transcript, and consequence of edits on coding capacity. Only transcripts with TPM ≥ 1 were considered in analysis of editing consequences. R statistical computation software with the following packages was used to parse VEP output: sleuth, biomaRt, dplyr, plyr, AnnotationFuncs, org.Mm.eg.db, ggplot2 [69–71].

### Validation of identified A-to-I RNA edits

Total RNA was isolated by TRIzol extraction of GV oocytes collected from wild-type PMSG stimulated C57B/6J female mice and treated with DNase I (Thermo Fisher, AM2222). cDNA was generated using Superscript III (Thermo Fisher, 18080044) and Oligo dT_20_ primers (Thermo Fisher, 18418020). Genomic DNA (gDNA) was isolated from tail clip of wild-type C57B/6J mice using REDExtract-N-Amp™ Tissue PCR Kit, (SigmaAldrich, XNAT-100RXN. gDNA and cDNA were amplified using Phusion® High-Fidelity DNA Polymerase (NEB, M0530). PCR reactions were purified using AMPure XP (Beckman Coulter, A63987) bead purification and cloned using the Zero Blunt™ TOPO™ PCR Cloning Kit (Thermo Fisher, K286020). Transformations were grown on LB+Kanamycin at 37°C overnight. Individual colonies were picked and grown in liquid culture, LB+Kanamycin, overnight at 37°C. Plasmid DNA was extracted from liquid culture using QIAprep Spin Miniprep Kit (Qiagen, 27104). Individual clones were Sanger sequencing using M13 Fwd and M13 Rev primers at GeneWiz (Boston, MA). All primers used for cloning and sequencing are listed in Supplemental Table 1.

### Western blot analysis

Western blots were performed on groups of 30 oocytes (PND12, GV, and MII eggs) isolated from untreated PND12 and from day 21 wild-type C57/B6J females stimulated with PMSG. MII eggs were *in vitro* matured by removal of milrinone (2.5 μM) for 16 hours. Eggs with visible polar bodies were used for analysis. Groups of 30 oocytes or eggs were lysed directly in SDS sample buffer with 5% β-mercaptoethanol, and subjected to electrophoresis using 10% SDS-PAGE in running buffer at a constant 120 V for 1 h. Proteins were electro-transferred onto PVDF membranes 350mA for 1 h at 4°C, and blocked with 5% (w/v) skim milk in Tris-buffered saline with 0.05% (v/v) Tween-20 (TBST) for 1 h at room temperature. Membranes were then probed with primary anti-ADAR1 (1:1000; sc-73408, Santa Cruz, Dallas, TX), or anti-Actin (1:5,000; sc-1616, Santa Cruz, Dallas, TX) antibody overnight at 4°C in TBST/BSA (50 mM Tris, 150 mM NaCl, 0.05% Tween20, 1.5% BSA) followed by incubation with secondary antibodies (mouse IgG HRP clone NA931V, GE Healthcare Life Sciences, Pittsburgh, PA; or goat IgG HRP, clone sc-2020 Santa Cruz) for 2 h at room temperature in blocking buffer. Membranes were washed three times in TBST and detected by enhanced chemiluminescence (Thermo Fisher Scientific, Waltham, MA).

### Statistics

Statistical analyses were performed using R and Prism 7.0. Results of multiple repeats were presented as means ± SEM. Bartlett’s tests were done to ensure equal variance among treatment groups. If data was normally distributed parametric tests were used. If data showed a variance outside of normal, the non-parametric test were used to determine if statistical differences existed. In cases where only two treatment groups existed, differences were determined by a non-parametric, unpaired t-Test (Mann Whitney). To determine statistical differences between groups with more than two treatment groups, one-way analysis of variance (ANOVA) followed by Tukey’s multiple comparison tests were used, or non-parametric Kruskal-Wallis test with Dunn’s multiple comparisons test were performed. In samples with two variables, a two-way ANOVA followed by Tukey’s multiple comparison tests was performed, X^2^ tests were used where appropriate to determine observed versus expected significance and to test differences in populations. Values of p < 0.05 were considered significant.

## List of abbreviations

ADAR: adenosine deaminases acting on double stranded RNA
*Adad2*: adenosine deaminase domain containing 2
BAM: Sequence Alignment Map (SAM)
CDS: coding region sequence
GATK: Genome Analysis Toolkit
GV: germinal vesicle oocyte
MII: metaphase II egg
LINE: long interspersed nuclear elements
ORF: open reading frame
PND12: postnatal day 12
SEM: standard error of the mean
SINE: short interspersed nuclear elements
TPM: transcripts per million

## Declarations

### Ethics approval and consent to participate

All animal experimentation has been approved by the University of Kansas Medical Center Institutional Animal Care and Use Committee.

### Consent for publication

Not Applicable

### Availability of data and material

RNA sequencing data for postnatal day 12 oocytes has been deposited under SRP133083.

### Authors’ contributions

PB, NSA, and LKC designed and PB, NSA, and XH performed experiments. KAV and KEL performed sequencing. PB, NSA, KEL, and LKC wrote and edited the manuscript.

## Acknowledgements

Not applicable

---

**Supplemental Figure 1. Adenosine deaminase genes are lowly expressed in mouse oocytes and eggs.** A) Principal components analysis on RNA-seq datasets that were utilized in our studies. *Adarb1* (B), *Adarb2* (C-E), and *Adad2* (F) isoform abundance in PND12 oocytes, GV oocytes, and MII eggs. TPM: transcripts per million. ^a,b^ Means ± SEM within a panel that have different superscripts were different (p < 0.05); Kruskal-Wallis test.

**Supplemental Figure 2. Schematic diagrams showing *Adar* variants.** A) *Adar* transcript variants are labeled according to NCBI Reference Sequence. Variant 3 and variant 1 generate the longest mRNA isoforms, containing 15 exons. Variant 1, however, has a truncated exon 7 that is the result of alternative splicing. Variant 2 utilizes an alternative promoter and alternative start codon in exon 2. B) Exon 7 sequences of the three *Adar* variants detailing the truncated region of variant 2. C) Protein domains of ADAR1 variants. Variant 1 is shorter than variant 3 by 26 amino acids that are missing between the dsRNA binding domain and the deaminase domain.

**Supplemental Figure 3. Patterns of RNA editing in somatic cells.** A) Proportion of the transcriptome that contains RNA edits within colon, heart, large intestine, stomach, brain, and brain from the ADAR1 E861A mutant mouse. B) The frequency of edited transcripts (y-axis) exhibiting one or multiple edits per transcript in somatic tissues. The x-axis is the sqrt (square root) normalized, while the numbers above peaks represent the actual number of edits per transcript. C) Total number of mRNA transcripts per sample. Only transcripts with TPM ≥ 1 were analyzed.

**Supplemental Figure 4. Adenosine deaminase expression in somatic tissue.** The expression of A) *Adar*, B) *Adarb1*, and C) *Adarb2* was determined for the following somatic tissue; Brain, Brain E861A, Colon, Heart, Stomach, and Large Intestine. TPM: transcripts per million.

**Supplemental Figure 5. Codon usage in PND12 oocytes, GV oocytes, and MII eggs.** The frequency of codons occurring in all transcripts with a TPM ≥ 1.

**Supplemental Table 1.**
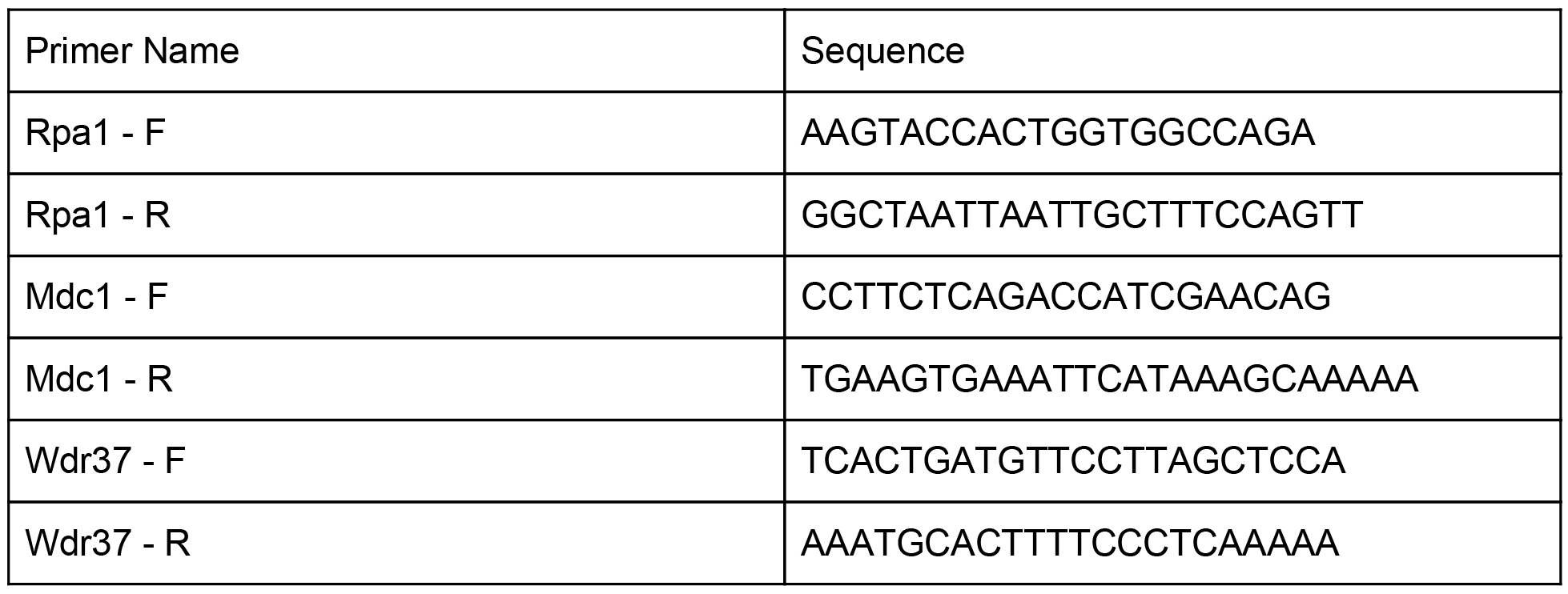
Primer sequences used to validate RNA edits.

